# Amino Acid Stress Induces Non-AUG Initiation of c-myc Translation

**DOI:** 10.64898/2026.06.01.729300

**Authors:** Miao Zhang, Changqian Zhou, Joao Paulo, Steven Gygi, Tracy Keller, Malcolm Whitman

## Abstract

Initiation of translation at non-AUG start codons can generate novel isoforms of important cellular regulators, with activities or localization distinct from their AUG-initiated counterparts. Ribosomes scanning the 5’UTR upstream of canonical AUG starts recognize near-cognate starts inefficiently, however, and the mechanisms by which non-canonical starts might be regulated physiologically are poorly understood. We show here that the restriction of utilization of individual amino acid induces a switch in expression of c-myc from its AUG-initiated isoform, c-myc-2, to an upstream CUG-initiated isoform, c-myc-1. This switch is enhanced in cells lacking GCN2, the protein kinase linking amino acid stress to phosphorylation and inhibition of eif2α. Induction of upstream CUG start site usage in c-myc by restriction of proline utilization depends on the proximity of proline codons downstream of the canonical AUG start site. These observations support a model in which the stalling of an elongating ribosome near a canonical start leads to the accumulation of a ribosome queue that facilitates initiation at an upstream non-canonical start site. We find similar effects associated with deprivation of amino acids other than proline, and isoform changes with ATF-3 as well as c-myc. These findings suggest a broad mechanism by which amino acid stress can regulate translation initiation site use to modify the N-terminal proteome.

## INTRODUCTION

In the standard model for the initiation of protein translation, the 43S pre-initiation complex (PIC, composed of the 40S ribosomal subunit, GTP-bound EIF2, charged methionyl tRNA, and additional initiation factors) is recruited to the 5’ end of capped mRNA bound by the EIF4F complex^1,2^. This assembled 48S PIC complex then scans the 5’ UTR until it encounters an AUG start codon, at which point base pairing of the methionyl-tRNA at the AUG initiates translation. Although examples of translation at non-canonical start codons (commonly CUG) have been known for decades, recent application of ribosome profiling has dramatically increased the identification of instances of the usage of non-canonical initiation sites that are upstream of and in frame with canonical AUG starts^3,4^. The regulated use of upstream non-canonical initiation sites has the potential to substantially broaden the functional translatome, and multiple examples of changes in protein function as a result of N-terminal extensions of translation have been identified^5^.

Remodeling of the translatome is fundamental to the altered biology of tumor cells, and both oncogene products and tumor suppressors can be regulated by the use of alternative translation initiation sites^5,6^. Global changes in sites of translation initiation have been proposed to be essential for tumorigenesis in squamous cell carcinoma^6^, and key regulators of tumorigenesis, including Sox2^5^, FGF^7,8^, VEGF^9,10^, WT1^11^, and PTEN^12,13^ have each been shown to have isoforms initiated from starts upstream of a canonical AUG, with radically altered cellular distribution or function when compared to the the canonically initiated isoform. c-myc was the first oncogene product for which isoform translation from a non-canonical upstream start site was established^14^. c-myc-1(p67) is translated from a CUG found 14 codons upstream from the AUG start that initiates c-myc-2 (p64). c-myc-2 is typically associated with highly proliferative cells, whereas c-myc-1 expression is induced when cells in culture reach confluence^15^. In Burkitt’s lymphoma, loss of intron 1 of c-myc results in expression of c-myc-2 but not c-myc-1, and this change has been proposed to be a component of the oncogenicity of c-myc in this tumor type^14^.

How translation is initiated at non-AUG start sites is controversial, and may involve at least two distinct mechanisms^16^. A partial mismatch between the methionyl tRNA of the canonical 48S PIC and a non-canonical start codon(e.g. CUG) can be permissive for translation, and initiation at non-canonical start codons that differ by a single base from AUG has been generally thought to result from mismatch recognition by methionyl-tRNA^16,17^. Initiation of translation at CUG starts by tRNA^Leu^, a cognate match for CUG, has also been reported^18,19^, however. This mechanism does not involve EIF2, but instead uses the alternative initiation factor EIF2A, which has been proposed to have a broad role in the enhanced utilization of upstream start sites during tumorigenesis^6^.The relative contributions of canonical Met-tRNA/eIF2 containing PICs and non-canonical initiation complexes to initiation at non-canonical start codons therefore remains an important question in the field^17^.

Physiological mechanisms of regulation of canonical versus upstream non-canonical start site use are likewise incompletely understood. Non-canonical start sites are typically used much less efficiently than canonical AUG starts^16,20^, and so additional regulatory mechanisms are likely necessary to allow non-canonical starts to compete with canonical ones. Conditions that increase the residence time of PICs at a non-canonical start codon provide one mechanism to enhance initiation at these sites. The stalling of elongating ribosomes, either at inefficiently translated codons^20,21^ or as a result of direct pharmacologic inhibition, has been suggested to lead to the accumulation of queues of 48S PICs behind the stalled ribosome. If this “ribosome queue” extends upstream to the position of the upstream start site, the sustained juxtaposition of the 48S PIC with the otherwise disfavored start codon allows effective initiation of translation^20^. Although this model is attractive, physiological conditions that drive a shift in the relative use of canonical start sites and upstream, in frame, non-canonical ones have not been established.

We report here that the restriction of the availability for translation of individual amino acids can drive a shift in the utilization of canonical versus upstream start sites on c-myc mRNA. This shift occurs only when a codon cognate to the restricted amino acid occurs just downstream of the canonical start site. This finding strongly favors a model in which the queueing of PICs behind ribosomes stalled just downstream of a canonical start leads to preferential utilization of an upstream start site. We also present evidence that this process is likely to use canonical, Met-tRNA-containing, 48S PICs to initiate upstream translation, and that this process likely controls upstream start site use in other important cellular regulatory proteins in addition to c-myc. These findings establish a mechanism linking amino acid limitation, ribosome stalling, and transcript selective remodeling of the N-terminal proteome.

## RESULTS

### AARS inhibition induces slow migrating forms of c-myc and ATF3

Restriction of amino acid availability is well established to induce a conserved set of stress-responsive transcription factors, including c-myc and ATF3^22,23^. While investigating the transcriptional changes following restriction of proline utilization by the prolyl tRNA synthetase (PARS) inhibitor halofuginone (HF), we observed that HF induced two distinct protein forms of the stress-responsive factors c-myc and ATF3. Induction of both c-myc and ATF3 by HF occurred in GCN2-null HCT116 cells and, strikingly, expression of the slower migrating form of each factor was strongly enhanced in cells lacking GCN2 (**Fig.1**). These slow migrating isoforms of c-myc and ATF3 appeared over 6-12 hours after HF treatment (**Fig.S1A**). Induction of these isoforms occurred at the lowest doses (5-10 nM) of HF at which any biological response (e.g. ATF4 induction) was detectable in wild type cells (**Fig.1**). Induction of both isoforms of c-myc and ATF3 was reversed by addition of excess proline (**Fig.S1b**), which competes with HF at the PARS active site^24^, confirming the specificity of HF action via PARS inhibition. Induction was recapitulated by the threonyl tRNA synthetase inhibitor borrelidin^25^ (**Fig.S1c**), confirming that aminoacyl tRNA synthetase (AARS) inhibition is responsible for the induction of the slow-migrating isoforms of both c-myc and ATF3. Enhanced induction of the slower migrating isoforms was also seen in cells lacking GCN1, a sensor of ribosome collisions that acts upstream of GCN2^26^ (**Fig.S1d**). These findings establish that AARS inhibition is sufficient to induce expression of two distinct isoforms of c-myc and ATF3, and that the absence of GCN2 results in preferential expression of the slower migrating isoforms of each factor.

**Fig 1.**
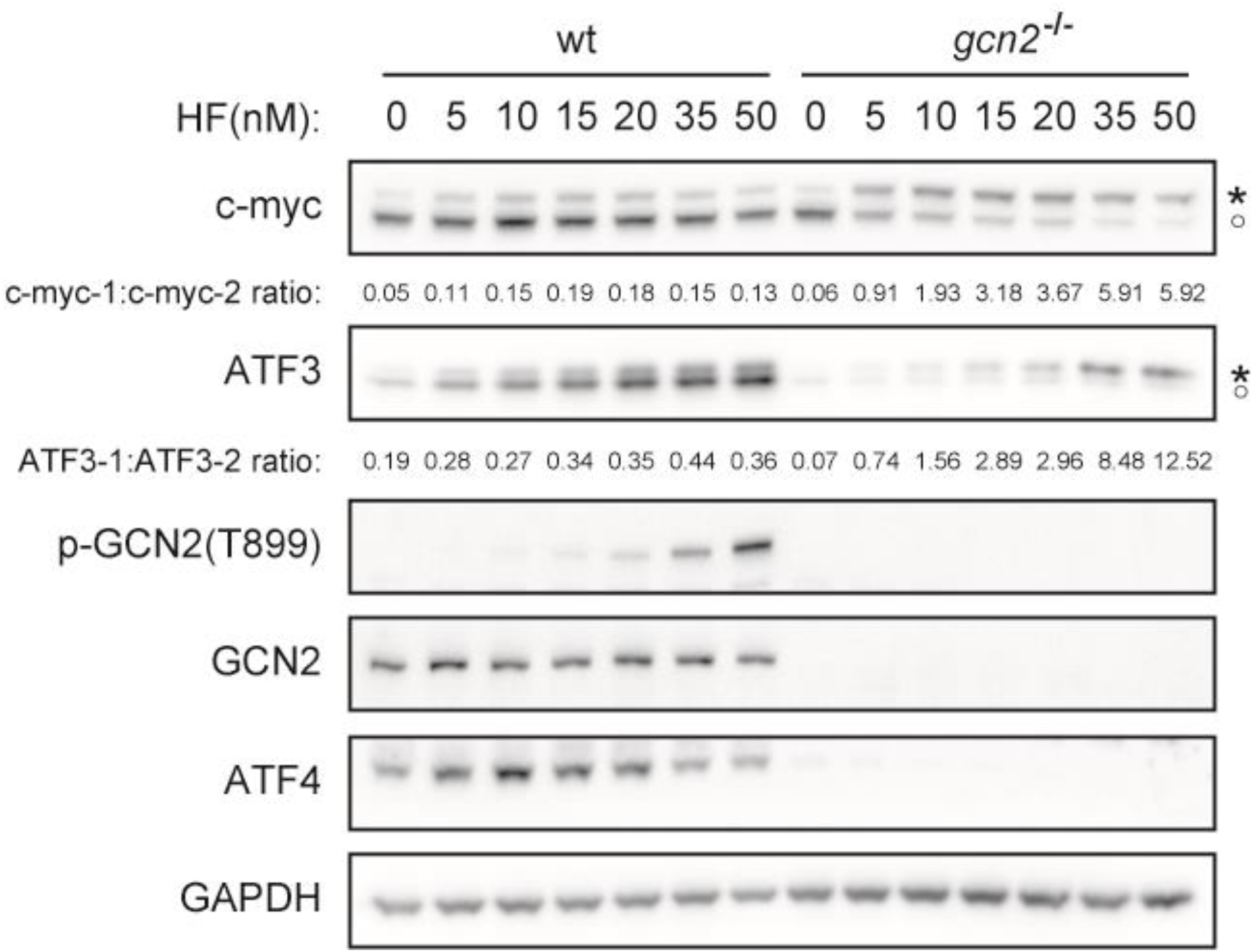
Amino acid stress induces slow migrating forms of c-myc and ATF3. HCT116^wt^ or HCT116(GCN2^-/-^) cells were treated with HF for 8 hours, lysed and analyzed by Western blot as indicated. Signal intensity for the isoforms of c-myc and ATF3 were determined using ImageJ analysis of the blots, and ratios of intensity for the slow migrating isoform versus fast migrating isoform of each protein are indicated. Asterisk, slow migrating isoform (c-myc-1, ATF3-1); black circle, faster migrating isoform (c-myc-2, ATF3-2).

### Changed migration of c-myc is due to an upstream CUG start to translation

Prior studies of c-myc have established that the use of alternate translation start sites can generate two isoforms of c-myc protein of sizes similar to those we observe in **Fig.1**^14^. The smaller, faster migrating, isoform (c-myc-2) results from initiation at a canonical AUG start, while the larger isoform (c-myc-1) is initiated at a non-canonical CUG start 14 codons upstream of the AUG start. Ectopic expression of HA-tagged c-myc translated from a canonical AUG start, but lacking the endogenous upstream CUG (**Fig.2A**), showed no change in migration following HF treatment (**Fig.2B**), pointing to alternative start site use, rather than a post-translational modification, as the likely basis for the change in c-myc size. Ectopic expression of c-myc using a construct that contain the endogenous 5’ UTR, including the upstream CUG, recapitulated regulation of endogenous c-myc migration, confirming this possibility (**Fig.2C**). Deletion of the majority of the 5’ UTR had no effect on this regulation, so long as the upstream CUG was retained (**Fig.S2**). Elimination of the canonical AUG start site resulted in expression of a c-myc isoform that co-migrated with the slower migrating, HF-induced, isoform, whereas elimination of the upstream CUG resulted in expression of the faster migrating constitutive isoform (**Fig.2D**). These findings indicate that the slow and fast-migrating forms of c-myc we observe correspond to c-myc-1 and c-myc-2, respectively.

**Fig.2.**
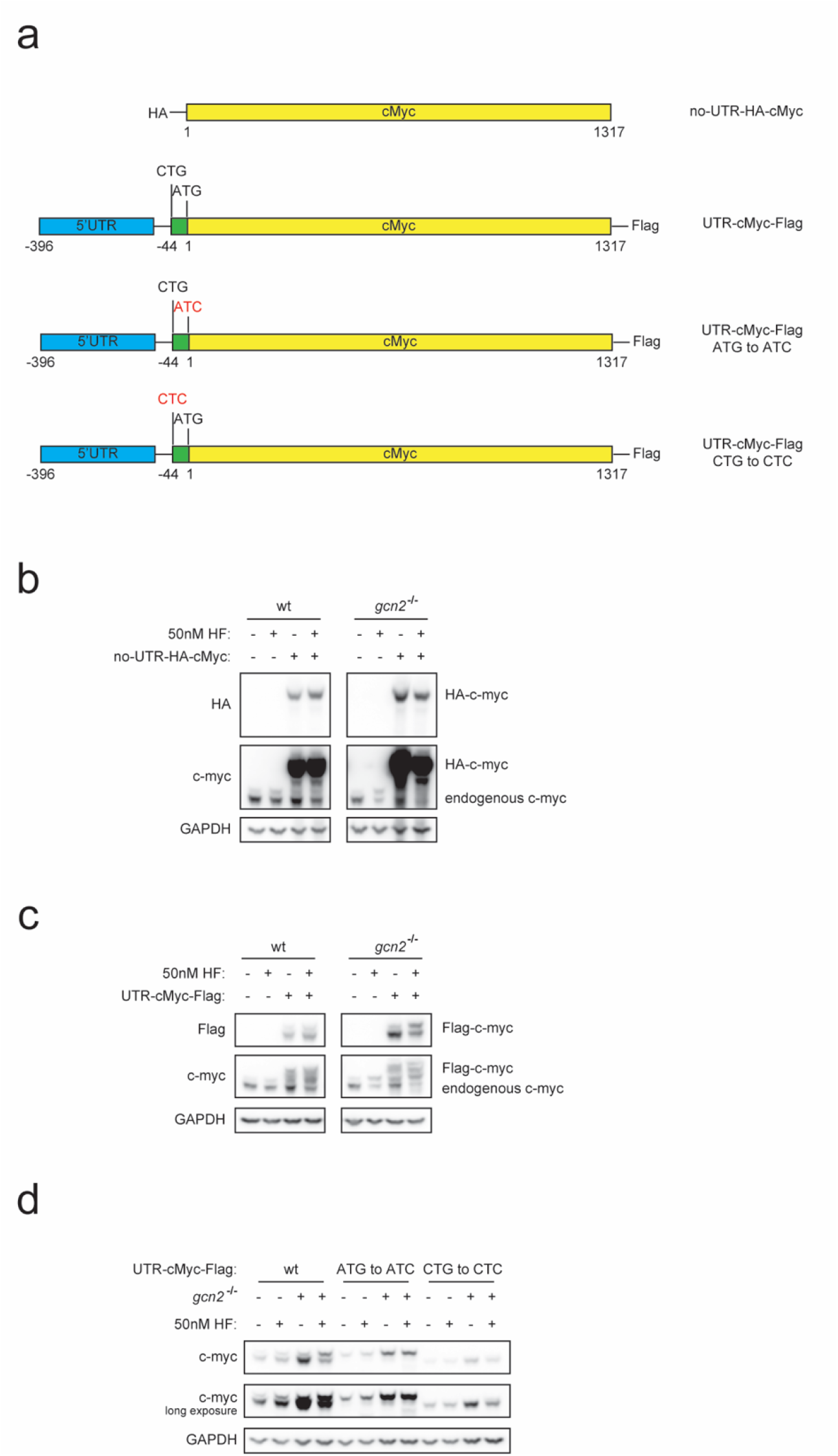
Altered migration of c-myc is due to an upstream CUG translational start. a) Scheme of c-Myc constructs designed to analyze 5’UTR function. (b) HCT116^wt^ or HCT116(GCN2^-/-^) cells were transfected with no-UTR-HA-cMyc for 24 hours, treated with HF for 8 hours and then analyzed by Western blot; c) HCT116^wt^ or HCT116(GCN2^-/-^) cells were transfected with UTR-cMyc-Flag and treated and analyzed as in (b); d) HCT116^wt^ or HCT116(GCN2^-/-^) cells were transfected with UTR-cMyc-Flag, UTR-cMyc^ATG to ATC^-Flag, or UTR-cMyc^CTG to CTC^-Flag, were treated and analyzed as in (b).

Initiation at CUG codons can result from mismatch recognition of the CUG by Met-tRNA as part of the canonical 48S pre-initiation complex (PIC)^16,27^. The amino acid encoded by the CUG start codon for c-myc-1 is, accordingly, annotated in Uniprot as methionine, but direct evidence for the use of methionine at this position is lacking. An alternative possibility for CUG initiation has been proposed by Starck et al^18^, who found evidence that tRNA^Leu^, in association with the non-canonical initiation factor eIF2A, is used to initiate translation at upstream CUG starts to generate antigenic peptides delivered to the MHC. This study used toeprinting analysis of c-myc mRNA to identify tRNA^Leu^ at the upstream CUG start. To establish whether HF-induced translation of c-myc-1 is initiated by tRNA^Met^ or tRNA^Leu^, we performed LC-MS/MS analysis of immuno-isolated c-myc-1 from HF treated cells (**Fig.S3A**). The predicted N-terminal peptide was found to contain only methionine at position 1, indicating that partial mismatch recognition of the CUG by tRNA^Met^ predominates for the initiation of translation of c-myc-1. We also used CRISPR inactivation of eIF2A to examine whether this non-canonical factor was important for translation of c-myc-1 (**Fig.S3B**). Reduction in eIF2A levels had no effect on HF-induced expression of Myc-2. These findings indicate that c-myc-1 translation at its CUG start in response to HF treatment is primarily initiated by a canonical tRNA^Met^-containing 48S PIC.

### Blocking inhibition of eIF2α recapitulates loss of GCN2

GCN2, like other ISR family kinases, regulates translation initiation through inhibition of eIF2α activity by direct phosphorylation^28^. To confirm that the loss of GCN2 enhanced c-myc-1 translation following HF treatment as a result of reduced inhibition of eIF2α, we treated wild type cells with HF in the presence of ISRIB, a small molecule that restores eIF2α activity by inhibiting eIF2b^29^. ISRIB enhanced induction of c-myc-1, indicating that the effects of loss of GCN2 are likely mediated through its regulation of eIF2α (**Fig.3**). HF treatment of cells in which eIF2α can no longer be regulated by ISR kinases (eIF2α(S51A)^30^) recapitulated the loss of GCN2 with respect to enhancing induction of c-myc-1, further supporting the likelihood that HF-stimulated phosphorylation of eIF2α by GCN2 limits c-myc-1 translation in wild type cells. Induction of eIF2α phosphorylation following ER stress with thapsigargin (through the ISR kinase PERK)^31^ did not induce c-myc-1, and had a weak but detectable inhibitory effect on c-myc-1 induction by HF relative to c-myc-2 induction (**Fig.S4**), also supporting the interpretation that increased eIF2α phosphorylation reduces translation from the non-canonical CUG start. Under each condition tested, induction of ATF3^slow^ tracked with induction of c-myc-1, suggesting that a common mechanism underlies the generation of alternative isoforms of both ATF3 and c-myc.

**Fig.3.**
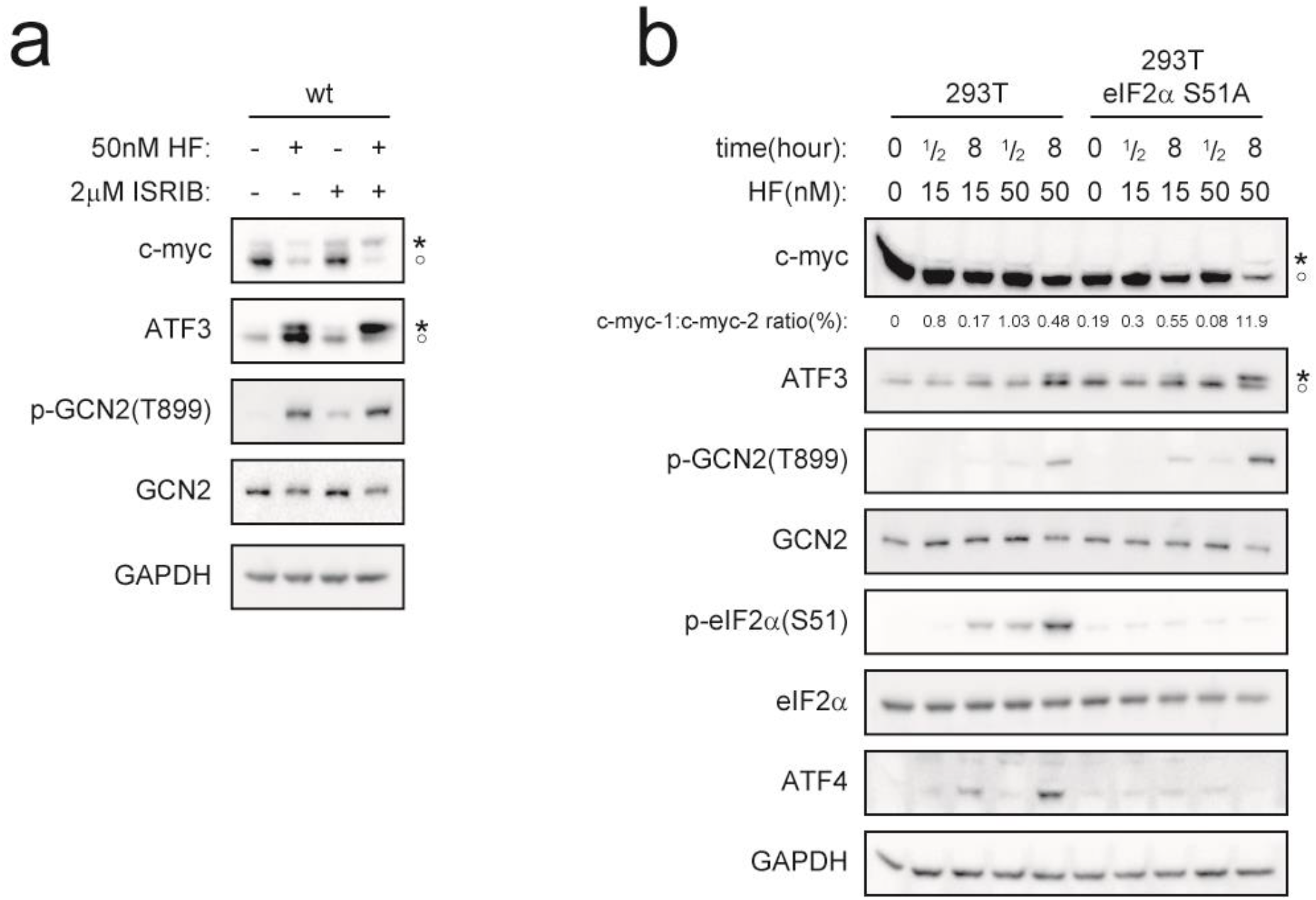
Regulation of eIF2α accounts for the effect of GCN2 loss on c-myc-1 induction. a) HCT116^wt^ cells were treated with vehicle control, HF, ISRIB, or HF+ISRIB for 8 hours, and analyzed by Western blot. b) 293T^WT^ or 293T(eIF2α^S51A^) cells were treated with 0-50 nM HF for indicated times and analyzed by Western blot. Band intensities were analysed by ImageJ and intensity ratios for c-myc-1 and c-myc-2 are indicated.

### Start-proximal codons determine effects of AARS inhibition on upstream CUG use

Little is known about how the transition from the usage of a canonical AUG start to an upstream non-canonical start is regulated. Cell treatments that slow the progress of translating ribosomes can enhance upstream CUG usage, leading to the proposal that the queuing of 48S PICs behind slowed or stalled ribosomes enables translational starts at otherwise disfavored non-canonical start sites ^20^. Limitation of amino acid availability or utilization is well established to stall translating ribosomes at codons cognate to the amino acid under restriction ^26,32-35^, providing a potential mechanism for codon-specific induction of 48S PIC queuing, and consequent upstream CUG initiation. To test this possibility, we examined the effect of mutation of AUG-proximal proline codons on the ability of HF to induce c-myc-1 (**Fig.4a,b**). Mutation of two AUG-proximal proline codons downstream of the AUG prevented c-myc-1 induction by HF, whereas mutation of proline codons upstream of the AUG had no effect. Mutation of proline codons had no effect on c-myc-1 induction by borrelidin **(Fig.S5**). These findings establish that proline codons occuring downstream of the canonical AUG start are critical for the enhancement of c-myc-1 expression in response to prolyl tRNA synthetase inhibition by HF.

**Fig 4.**
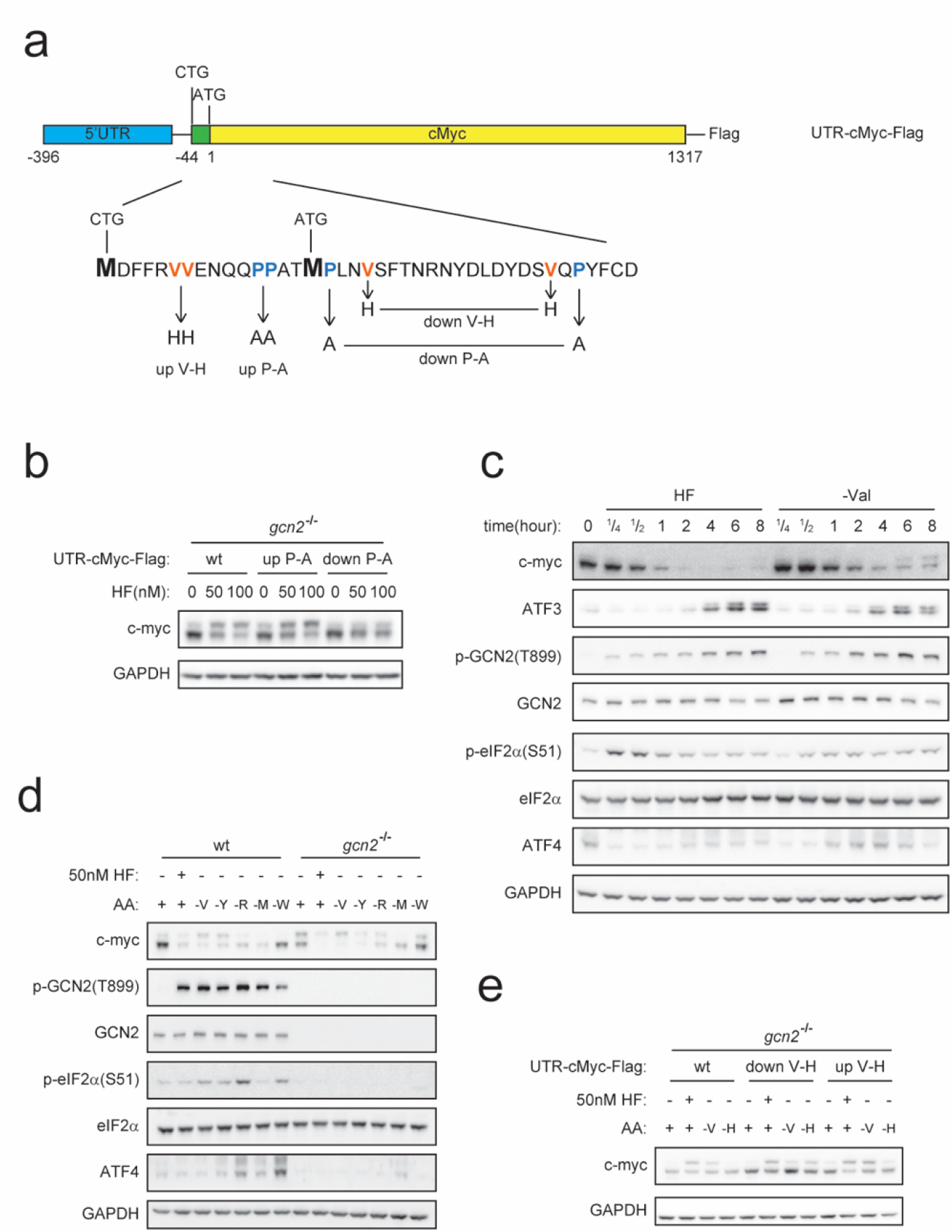
AUG start-proximal codons determine upstream CUG use under amino acid stress. a) Scheme of c-Myc constructs designed to test the role of cognate codons in cMyc-1 induction. Codons indicated in red (valine) or blue (proline) were mutated to histidine or alanine as indicated. “up-P-A” denotes proline codons mutated to alanine upstream of the canonical AUG start, “down-P-A’ codons mutated downstream of the AUG; “up V-H” and “down V-H” likewise denote the position of mutated codons relative to the AUG start. b) HCT116(GCN2^-/-^) cells were transfected with UTR-cMyc-Flag, UTR-cMyc-Flag^up P-A^, or UTR-cMyc-Flag^down P-A^. Then treated with HF for 8 hours,and analyzed by Western blot. c) HCT116^wt^ cells were incubated with HF in DMEM or with DMEM lacking valine for indicated times. d) HCT116^wt^ or HCT116(GCN2^-/-^) cells treated with HF, or incubated with control DMEM medium or DMEM lacking one one of the amino acids valine, tyrosine, arginine, methionine, or tryptophan for 8 hours. e) HCT116(GCN2^-/-^) cells were transfected with UTR-cMyc-Flag, UTR-cMyc-Flag^up V-H^, UTR-cMyc-Flag^down V-H^, and then treated with HF in standard DMEM, or incubated with DMEM medium without valine, or histidine for 8 hours.

c-myc-1 and ATF3^slow^ were also potently induced following removal of valine from culture medium (**Fig.4c**), establishing that amino acid deprivation recapitulates the effects of treatment with AARS inhibitors. In contrast to the effects of HF and borrelidin, however, preferential induction of c-myc-1 and ATF3^slow^ in response to valine deprivation was indistinguishible in wild type and GCN2^-/-^ cells (**Fig. 4d**). To examine more broadly the effects of amino acid deprivation on c-myc-1 induction, we tested the effects of starvation of cells for each of several amino acids on c-myc-1 induction (**Fig.4d**). Depletion of each amino acid tested activated GCN2 phosphorylation, confirming that in each case amino acid limitation had become limiting for tRNA charging, but depletion efficacy varied widely with respect to induction of c-myc-1. Depletion of different amino acids had variable effects on the induction of ATF4, which may reflect additional regulatory mechanisms that complement effects on eIF2α activity on ATF4 translation. To exclude the possibility that ATF4 induction may modify the induction of c-myc-1 and ATF3^slow^, we established that neither ectopic expression or CRISPR inactivation of ATF4 had any effect on induction of c-myc-1 and ATF3^slow^ (**Fig.S5C**,**D**). These results indicate that conditions that activate GCN2 phosphorylation are not alone sufficient for c-myc-1 induction, and that variability in ATF4 induction does not account for the variable efficacy in c-myc-1 induction by restriction of different amino acids.

To further investigate the possibility that codon position relative to the AUG start is important for induction of upstream translational starts, we examined the effects of altering AUG-proximal codons on c-myc-1 induction by amino acid deprivation. Valine has two cognate codons within 20 codons downstream of the canonical AUG start of c-myc, and valine deprivation potently induces c-myc-1 expression. In contrast, tryptophan and histidine do not have AUG-proximal codons (the first tryptophan codon occurs at +50 with respect to the canonical AUG, first histidine codon at +168), and neither tryptophan nor histidine deprivation increased c-myc-1 expression (**Fig.4d,e**), suggesting that proximity of cognate codons to the AUG start is critical for amino acid deprivation driven upstream start usage. To establish a causal role for AUG-proximal codons in c-myc-1 induction by amino acid deprivation, we mutated the two AUG-proximal valine codons to histidine codons **(Fig.4e**). In this codon-switched construct, induction of c-myc-1 by valine deprivation was reduced, but induction by histidine deprivation was enhanced, and induction by HF was unaffected. The presence of AUG-proximal codons cognate to a restricted amino acid are therefore both necessary and sufficient for the induction of upstream start usage on c-myc mRNA by amino acid restriction.

The novel HF-induced isoform of ATF3 is likewise induced by deprivation of amino acids with AUG-proximal codons (**Fig.S5b**). The tumor suppressor PTEN has several functionally distinct isoforms generated by alternative start site use ^12,13^. Canonically initiated PTEN migrates as an ∼50 kD protein, whereas the principle CUG-initiated isoform, PTENα, migrates as a 65-70 kD protein^12^. A PTEN isoform corresponding to the size of PTENα was induced by deprivation of arginine, with 3 AUG-proximal cognate codons, and very weakly by deprivation of valine, with one AUG proximal cognate codon (**Fig.S5d**). Collectively, these findings establish the codon-specific regulation of translation start site choice by the inhibition of amino acid availability or utilization, and suggest that this may be a widespread phenomenon for the modification of important regulators of cell function and phenotype.

## DISCUSSION

The regulated metabolism of amino acids in tissues is increasingly recognized to have multiple roles in tissue homeostasis and pathology, including the regulation of local inflammatory, fibrotic, and cell death responses, as well as the modulation of a broad range of both innate and adaptive immune responses ^36-38^. The activation of GCN2 is key to many of these effects, but GCN2-independent effects of local amino acid limitation on tissue biology have also been identified^39^. We report here that both AARS inhibitors and amino acid restriction can modify the N-terminal proteome by driving the enhanced use of non-canonical upstream start sites. In contrast to previously described triggers of upstream start site usage, amino acid restriction requires the presence of codon(s) cognate to the restricted amino acid just downstream of the canonical start site to induce upstream start usage. These findings imply that depletion of amino acids in tissues may have effects on mRNAs with non-canonical start sites that differ depending on the amino acid that is depleted. The local insufficiency of individual amino acids can occur as a result of the local induction of endogenous amino acid degrading enzymes (e.g. degradation of tryptophan by cytokine-induced IDO1)^37^, the therapeutic application of exogenous amino acid degrading enzymes (e.g. asparaginase for leukemia)^40^, or elevated amino acid utilization in tissues (e.g. arginine metabolism for nitric oxide production)^41^. In addition, systemic availability of amino acids can be limiting for tissue specific metabolic needs (e.g. branched chain amino acids utilization in muscle). The selective modification of the N-terminal translatome by changes in the availability of individual amino acids therefore constitutes a novel mechanism for tissue regulation in a broad range of physiological and pathological circumstances.

Several lines of evidence presented here point to the induction of “ribosome queues” as a result of stalling at codons just downstream of the canonical AUG start as the basis for the enhanced utilization of the upstream CUG start on c-myc mRNA. In this model, queues of 48S PICs accumulate upstream of ribosomes that have stalled, either as a result of general inhibition of elongation (e.g. by cycloheximide)^20^ or as a result of specific features of the translated mRNA^21^. If this queue extends upstream of the stall site to an otherwise disfavored non-canonical start site, the prolonged positioning of a 48S PIC at the non-canonical site becomes permissive for the initiation of translation. The efficiency of this process will depend on the rate at which 48S PICs accumulate relative to the rate of resolution of the ribosomal stall, and on how many 48S PICs must accumulate to reach an upstream start site. Consistent with this model, we find that mutation of codons cognate to a limiting charged tRNA (e.g. Pro-tRNA^Pro^ following HF treatment) just downstream of the canonical AUG start in c-myc is sufficient to prevent upstream start utilization. The c-myc coding sequence contains 33 proline codons in addition to the two AUG-proximal codons mutated in Fig.4., indicating that proximity to the translation start site, rather than total codon number, is what determines the propensity of 48S PICs to accumulate behind the site of a ribosome stall. At codons positioned further downstream of the start site, fully assembled, elongating 80S ribosomes would collide with ribosomes stalled at proline codons, generating ribosome collisions that would be resolved by the RQC apparatus, thereby preventing accumulation of queues upstream of the collision. c-myc-1 induction has also been reported to occur as a result of deprivation of glutamine^42^, for which a codon occurs at AUG+34, and 5 consecutive codons at +48. Whether extended stretches of codons for a particular amino acid are important for queue induction will also be of interest or further investigation.

In these studies, we found that restriction of methionine did not alter the distribution of c-myc-1 and c-myc-2 (**Fig.4**), which is not surprising since our data suggest that methionine is used to initiate translation at both canonical and non-canonical start sites in c-myc, and the next methionine in the c-myc sequence occurs 100 codons downstream of the canonical AUG start. A previous study, however, reported induction of c-myc-1 by methionine depletion in confluent cells^15^. Whether this difference is associated with confluence, different cell types examined, or other factors, will require additional study.

The ribosome queueing model outlined above carries several additional implications regarding how upstream start site usage is controlled. The rate of accumulation of 48S PICs upstream of a stall site will be constrained by the availability of assembled 48S PICs. Activation of the GCN2 or other ISR kinases, which phosphorylate eIF2α to limit active 48S PIC assembly would therefore be expected to reduce the efficiency of upstream start use. This implication would explain our observation that the induction of c-myc-1 by HF is strongly enhanced in GCN2^-/-^ cells, or by reversal of the effects of eIF2α phosphorylation with ISRIB. Why GCN2 inactivation has less impact on c-myc-1 induction by valine deprivation in comparison to HF treatment, however, is less clear. It has been suggested that the perdurance of ribosome stalls may differ depending on the charged tRNA that is unavailable^35^, and the perdurance of a stall may impact the sensitivity of upstream start site use to 48S PIC availability. The distance between a canonical start site and an upstream start is also likely to be a critical variable in the effect of amino acid restriction on upstream start usage, since greater distance requires that a greater number of 48S PICs accumulate to reach the upstream start codon. In the case of c-myc, the CUG start is only 14 codons upstream of the canonical start site. In PTEN, in contrast, the CUG start initiating PTENα is 173 codons upstream of the canonical start, potentially requiring much longer perdurance of stalls to provide time for the accumulation of 48S PICs sufficient for upstream start site engagement. Arginine deprivation, which we find induces a long isoform of PTEN (**Fig.S5**), has been reported to induce long-lived ribosome stalls, potentially accounting for its efficacy in the induction of PTEN^long^. Further study of a range of mRNAs with inducible upstream start sites will be necessary to define the parameters that define the efficacy of upstream start induction by different amino acids.

C-myc has a central role in the regulation of cell proliferation, tumorigenesis, and stem cell establishment and maintenance^43-46^. The induction of a functionally distinct isoform of c-myc by amino acid restriction may therefore have a broad range of physiological implications. Although early studies found similar activities for c-myc-1 and c-myc-2 in the co-transformation of cells with bcr-abl^47^, subsequent work has identified potentially important differences in the functions of c-myc-1 and c-myc-2. C-myc-1 over-expression inhibits cell proliferation, in contrast to the proliferation promoting effects of c-myc-2^48^. Over-expression of each isoform induces substantially different patterns of gene expression^48,49^, and c-myc-1 interacts with a promoter element not recognized by c-myc-2^49^. It is also potentially important that c-myc is an important regulator of amino acid metabolism^44^, and c-myc-1 and c-myc-2 differ with respect to induction of known regulators of amino acid metabolism^48^, suggesting a possible regulatory feedback loop linking amino acid metabolism and c-myc isoform expression. Although these differences are intriguing, how the addition of 14 amino acids at the N-terminus, a largely unstructured region distal to the known functional domains of c-myc, might change the transcriptional activity of c-myc is not understood. This remains an important area for future investigation.

In this work we noted a change in migration of a second stress-responsive transcription factor, ATF3, in response to HF treatment that closely paralleled the change in isoform expression we observed for c-myc. LC-MS/MS analysis of ATF3 from HF treated cells did not yield data regarding N-terminal peptides (not shown), and we have not otherwise undertaken to characterize the slower migrating form of ATF3 induced by HF or amino acid restriction. The 5’UTR of ATF3 contains a CUG in a strong Kozak context 10 amino acids upstream, and in frame, with the canonical AUG start, consistent with the size of the novel ATF3 isoform we observe. Strikingly, the predicted upstream sequence is 90% conserved in ATF3 across a broad range of vertebrate species. The parallel regulation of c-myc and ATF3 isoforms suggests a broader regulatory motif linking amino acid restriction to changes in stress responsive transcription factors, but will require further characterization of the basis for the size change in ATF3 as well as its functional significance.

Regulation of start site utilization in response to amino acid stress provides a new mechanism for expanding the proteome in response to changes in the cellular environment. Alterations in start site utilization have drawn particular attention in the context of oncogenesis^5,6^, but the local regulation of amino acid availability has emerged as a key regulatory node in the regulation innate immunity and tissue homeostasis as well^36-38^. Investigation of the scope and functional effects of changes in the N-terminome as a result of amino acid restriction may therefore be of fundamental importance in understanding how local changes in amino acid availability modify tissue biology.

## MATERIALS AND METHODS

A complete list of all antibodies, reagents, cell lines, plasmids and oligonucleotides used in this study are listed in Tables S1-5, respectively.

### 1. Cell Culture and Analysis

#### 1.1 Cell Culture and amino acid starvation

HCT116 cells used in this study were purchased from ATCC and routinely tested for mycoplasma contamination. HEK293T eIF2αS51A cells and parental HEK293T cells were a kind gift of Dr. Laura Ranum, University of Florida. Cells were cultured in high glucose Dulbecco’s Modified Eagle Medium (DMEM) supplemented with 10% fetal bovine serum (Gibco), 100 U/mL penicillin G, 100 μg/mL streptomycin sulfate, and 250 ng/mL amphotericin B at 37℃ with 5% CO_2_. For amino acid starvation, cells were washed by PBS and cultured in DMEM lacking the indicated animo acid.

#### 1.2 Generation of Cell lines

For CRISPR-Cas9 mediated gene knockout, sgRNAs use were:

GCN1 (target sequence: AGACAC-TAAAGCGTTTTGCAGGG)

GCN2 (target sequence: ATGTCCCCCTTCGACCAGT)

ATF4 (#2 target sequence: GCCGATTCAGTTCTGCTTACCGG; #3 target sequence: AGTCCCTCCAACAACAGCAA)

eIF2A (#2 target sequence: ATTGACATAGGTGTCCATCC; #3 target sequence: AAGAGTTTCATCTTCTGACC) were cloned into the lentiCRISPRv2 vector (Addgene).

HCT116 cells were transfected using PEI MAX and selected with 2 μg/mL puromycin for 4 days. Following selection, cells were single-cell sorted by Flow Cytometry into 96-well plates. Individual cell clones were subsequently picked and identified.

#### 1.3 Plasmids and transient transfection

The pCSX constructs contain a 3xFlag tag at the 3’ end of the cMyc coding region. UTR-cMyc wt or mutants fragments were synthesized by gBlock (IDT) and inserted in the BamHI and NcoI site in the pCS-myc construct. Plasmid transfection was performed using PEI MAX (Polysciences) according to the manufacturer’s instruction.

#### 1.4 Western Blotting

Cells were washed twice with ice-cold PBS buffer and lysed in RIPA lysis buffer (50 mM Tris-HCl pH 7.4, 150 mM NaCl, 1% Triton X-100, 1% sodium deoxycholate, 0.1% SDS, 1 mM EDTA, 1x Protease Inhibitor Cocktail, and 1x PhosSTOP). The cell lysate was then clarified by centrifugation at 13,000 rpm at 4℃ for 10 minutes. Total protein concentration was measured using the BCA method (ThermoFisher Scientific) and normalized. Samples were supplemented with 1x Laemmli buffer and heated at 65℃ for 5 minutes.

Samples were loaded onto 4-12% Bis-Tris gels (ThermoFisher Scientific) and transferred to 0.2 μm nitrocellulose membranes. Membranes were blocked in 5% non-fat milk in TBST for 1 hour at room temperature and incubated overnight at 4℃ with the primary antibody diluted in blocking buffer. Membranes were washed with TBST (3x 10 minutes) and incubated with secondary antibody diluted in blocking buffer at room temperature for 1 hour. Membranes were then washed with TBST (3x 10 minutes) and imaged using a Syngene system. The band intensity of western blot was analyzed using Image J software. To ensure reproducibility, at least two fully independent experiments were conducted with consistent results.

### 2. LC-MS/MS Methods

#### Sample preparation for mass spectrometry

The precipitated samples were resuspended in 200 mM EPPS, pH 8.5 and digested at room temperature for 14 h with LysC protease at a 100:1 protein-to-protease ratio. Trypsin was then added at a 100:1 protein-to-protease ratio and the reaction was incubated for 6 h at 37°C. The digestion reaction was quenched with a final formic acid concentration of 2.5% and then desalted via StageTip, dried again via vacuum centrifugation, and reconstituted in 5% acetonitrile, 5% formic acid for LC-MS/MS processing.

#### Mass spectrometric data collection

Mass spectrometry data were collected using a Orbitrap Astral or Ascend mass spectrometer (Thermo Fisher Scientific, San Jose, CA) coupled with Neo Vanquish liquid chromatograph.

Using the Orbitrap Astral, peptides were separated using a 35-min gradient of 5 to 29% acetonitrile in 0.125% formic acid with a flow rate of 450 nL/min on a 50 cm uPAC column. The spray voltage was set at 2200 V. The scan sequence began with an Orbitrap MS1 spectrum with the following parameters: resolution 120000, scan range 350-1350 Th, automatic gain control (AGC) target 1,000,000, maximum injection time 50 ms, and centroid spectrum data type. We used a cycle time of 1 s for MS2 analysis which consisted of HCD (high-energy collisional dissociation) with the following parameters: AGC 20,000, maximum injection time 15 ms, isolation window 1.2 Th, normalized collision energy (NCE) 27%, and centroid spectrum data type. Dynamic exclusion was set to automatic. The FAIMS compensation voltages (CV) were -25, -35, -45, -55, -65V.

Using the Orbitrap Ascend, peptides were separated using a 90-min gradient of 5 to 29% acetonitrile in 0.125% formic acid with a flow rate of 450 nL/min on a 35 cm Accucore150 column. The spray voltage was set at 2800 V. The scan sequence began with an Orbitrap MS1 spectrum with the following parameters: resolution 60000, scan range 350-1350 Th, automatic gain control (AGC), target 100%, maximum injection time 123 ms, and centroid spectrum data type. We used a cycle time of 1 s for MS2 analysis which consisted of CID (collision-induced dissociation) with the following parameters: resolution 15000, AGC 150%, maximum injection time 27 ms, isolation window 1.2 Th, collision energy 30%, and centroid spectrum data type. Dynamic exclusion was set to automatic. The FAIMS compensation voltages (CV) were -30, -50, -70V for the first set and -45 and -60V for the second set.

#### Mass spectrometric data analysis

Mass spectra were processed using a Comet-based in-house software pipeline. MS spectra were converted to mzXML using MSconvert. Database searching included the protein of interest (cMyc) plus common contaminants which were concatenated with a reverse database composed of all these protein sequences in reversed order. The digest was set to non-specific. Searches were performed using a 3 Da precursor ion tolerance. Product ion tolerance was set to 0.02 Th. Oxidation of methionine residues (+15.9949 Da) and N-terminal acetylation (+42.0105 Da) were set as a variable modification. Peptide-spectrum matches (PSM) filtering was performed manually considering only tryptic peptides, an XCorr>1.5, and an absolute value of PPM mass tolerance <10. Spectra were manually validated.

