## Supplemental figures and tables for "Amino Acid Stress Induces Non-AUG Initiation of c-myc Translation"

Table S1.
Antibodies used in this study.

| Name | Source | ID |
|---|---|---|
| alpha Tubulin | GeneTex | Cat# GTX628802 |
| ATF3 | Santa Cruz Biotechnology | Cat# sc-188 |
| ATF4 | Proteintech | Cat# 10835-1-AP |
| cMyc | Cell Signaling Technology | Cat# 5605 |
| eIF2 alpha | Cell Signaling Technology | Cat# 9722 |
| eIF2A | Thermo Fisher Scientific | Cat# MA5-42514 |
| FLAG tag | Sigma-Aldrich | Cat# A8592 |
| GAPDH | Thermo Fisher Scientific | Cat# MA5-37687 |
| GCN1L1 | Bethyl | Cat# A301-843A |
| GCN2 | Santa Cruz Biotechnology | Cat# sc-374609 |
| GCN2 | Cell Signaling Technology | Cat# 3302 |
| HA tag | Sigma-Aldrich | Cat# 12013819001 |
| Phospho-eIF2alpha (Ser51) | Cell Signaling Technology | Cat# 3398 |
| Phospho-GCN2 (Thr899) | Thermo Fisher Scientific | Cat# PA5-105886 |

**Table S2.**

Chemicals and Reagents used in this study

| Name | Source | ID |
|---|---|---|
| Borrelidin | Sigma-Aldrich | Cat# B1936 |
| cOmplete™, EDTA-free Protease Inhibitor Cocktail | Roche | Cat# 5056489001 |
| GCN2 inhibitor A-92 | Axon Medchem | Cat# 2720 |
| Halofuginone | Selleck Chemicals | Cat# S8144 |
| PEI MAX | Polysciences | Cat# 24765-100 |
| PhosSTOP | Roche | Cat# 4906837001 |
| ISRIB | Sigma | Cat# SML0843 |
| Thapsigargin | Tocris | Cat#1138 |
| DMEM-amino acids | US Biological | Cat#D9800-27 |

**Table S3.**
Cell lines used in this study

| **Name** | **Source** | **ID** |
|---|---|---|
| HCT116 | ATCC | CCL-247 |
| HCT116 GCN1 KO | This paper | N/A |
| HCT116 GCN2 KO | This paper | N/A |
| HEK293T | Laura Ranum UFL | N/A |
| HEK293T eIF2α S51A | Laura Ranum UFL | N/A |

**Table S4.**
Plasmids used in this study

| Name | Source | ID |
|---|---|---|
| pCSX-6XHA-cMyc | This paper | N/A |
| pCSX-UTR-cMyc-3XFlag | This paper | N/A |
| pCSX-UTR-cMyc-ATG-ATC-3XFlag | This paper | N/A |
| pCSX-UTR-cMyc-CTG-CTC-3XFlag | This paper | N/A |
| pCSX-UTR-cMyc-P-A-3XFlag | This paper | N/A |
| pCSX-UTR-cMyc-V-H-3XFlag | This paper | N/A |
| pRk-ATF4 | Addgene | Addgene plasmid #26114 |
| pCSX-kozak-cMyc-3XFlag | This paper | N/A |

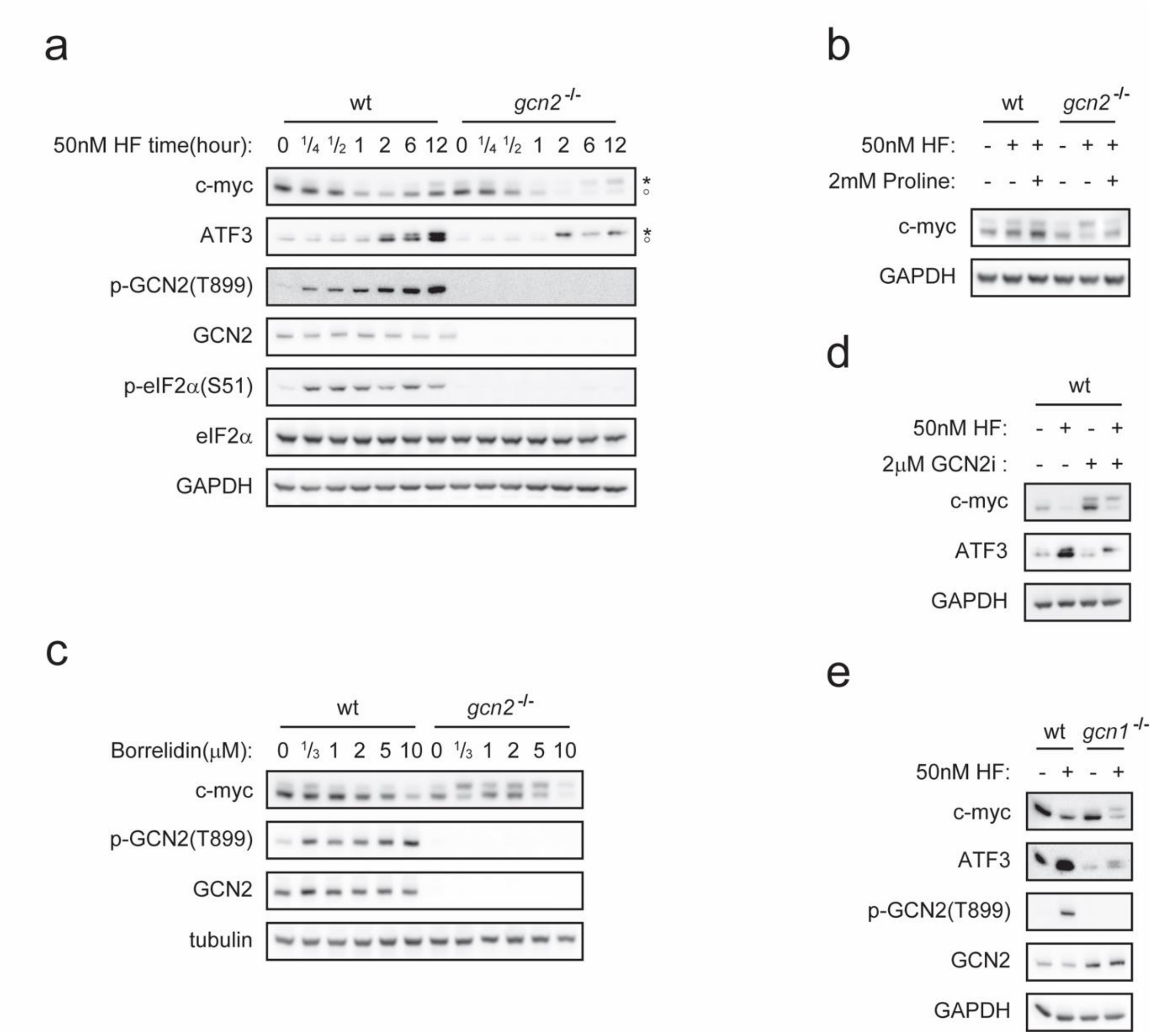


**Fig.S1**. **AARS inhibition induces long forms of c-myc and ATF-3 independent of GCN1 or GCN2.** a) HCT116$^{wt}$ or HCT116(GCN2$^{-/-}$) cells were treated with HF for times indicated and analyzed by Western blot. b) HCT116$^{wt}$ or HCT116(GCN2$^{-/-}$) HCT116 cells were treated with HF in the presence or absence of proline for 8 hours . c) HCT116$^{wt}$ or HCT116(GCN2$^{-/-}$) cells were treated with Borrelidin for 8 hours. d) HCT116$^{wt}$ cells were treated with HF in the presence or absence GCN2i for 5 hours. e) HCT116$^{wt}$ or HCT116(GCN2$^{-/-}$) cells were treated with HF for 8 hours.

**A**

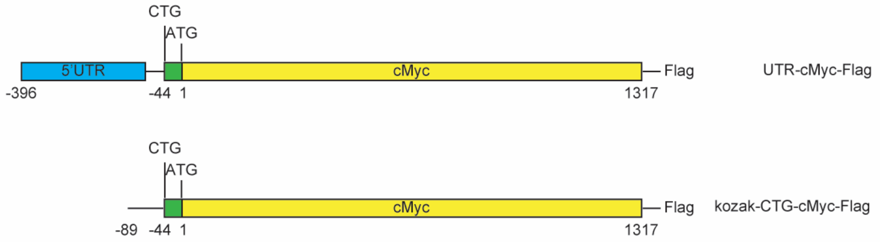


**B**

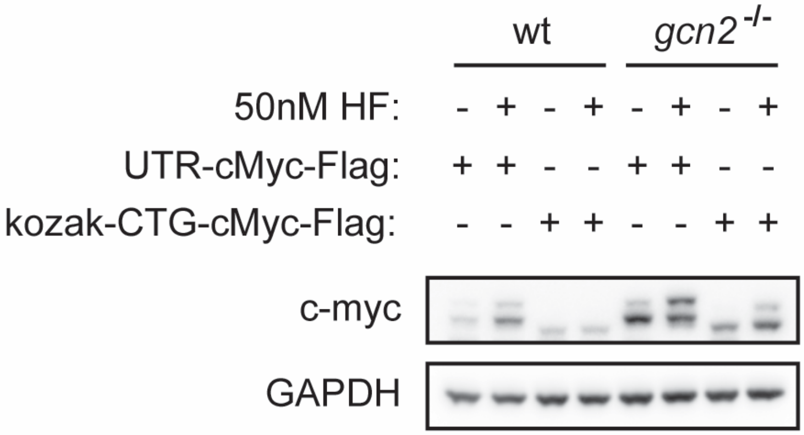


**Fig.S2.Translation of c-myc-1 from a minimal 5' UTR** A) Left panel: schematic of constructs use to test c-myc 5' UTR function. UTR-c-Myc-Flag contains full 5' UTR, kozak-CTG-c-Myc-Flag contains only a minimal Kozak consensus site upstream of the non-canonical CTG start site. B: HCT116$^{wt}$ or HCT116(GCN2$^{-/-}$) cells were transfected with UTR-c-Myc-Flag or kozak-CTG-c-Myc-Flag, then treated with HF for 8 hours.

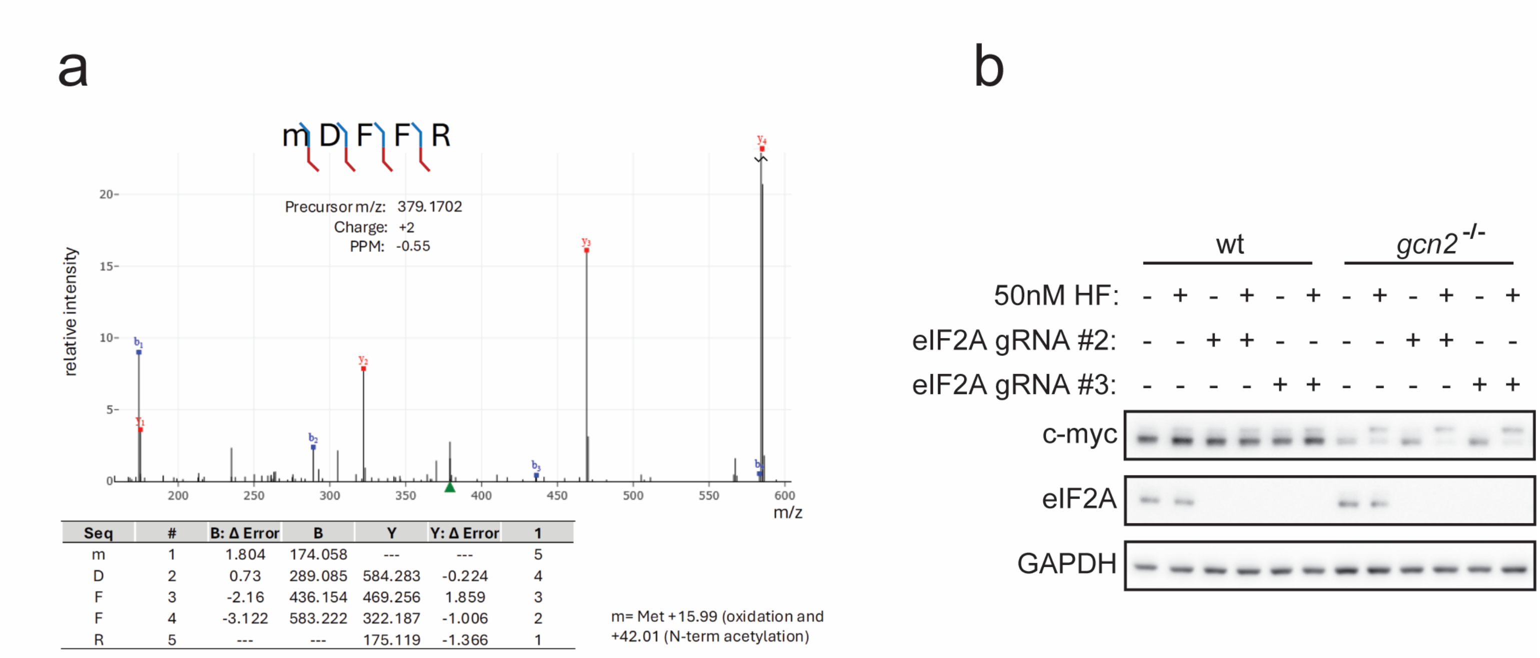


| Seq | # | B: Δ Error | B | Y | Y: Δ Error | 1 |
|---|---|---|---|---|---|---|
| m | 1 | 1.804 | 174.058 | --- | --- | 5 |
| D | 2 | 0.73 | 289.085 | 584.283 | -0.224 | 4 |
| F | 3 | -2.16 | 436.154 | 469.256 | 1.859 | 3 |
| F | 4 | -3.122 | 583.222 | 322.187 | -1.006 | 2 |
| R | 5 | --- | --- | 175.119 | -1.366 | 1 |

**Fig.S3**. **c-myc-1 CUG-intiated translation is likely to be initiated by a canonical 48S PIC**. a). Annotated MS/MS spectrum of the tryptic peptide mDFFR.The tandem mass spectrum corresponds to the doubly charged precursor ion (m/z 379.1702, +2 charge, −0.55 PPM error). Fragmentation produced a series of b- and y-ions that support the sequence mDFFR. The N-terminal residue (m) represents methionine containing two modifications: oxidation (+15.99 Da) and N-terminal acetylation (+42.01 Da). Fragment ions are annotated in red (y-ions) and blue (b-ions). The table below the spectrum lists observed b- and y-ion masses and mass errors for each residue position. b) Effect of EIF2A knockout on cMyc-1 synthesis in response to HF. Two independent sets of HCT116$^{wt}$ or HCT116(GCN2$^{-/-}$) lines in which EIF2A was inactivated by CRISPR (EIF2A gRNA#2; EIF2A gRNA#3) lines were generated and tested for HF induction of cMyc-1.

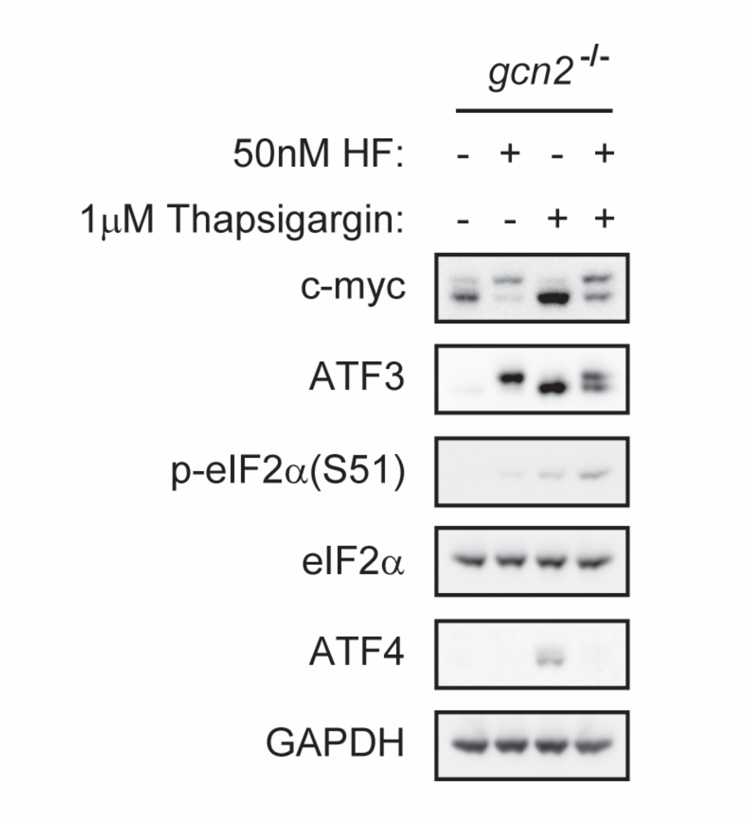


**Fig.S4**. **ER stress does not induce c-myc-1**.

HCT116(GCN2$^{-/-}$)cells were treated with HF or Thapsigargin for 8 hours, and analyzed by Western blot.

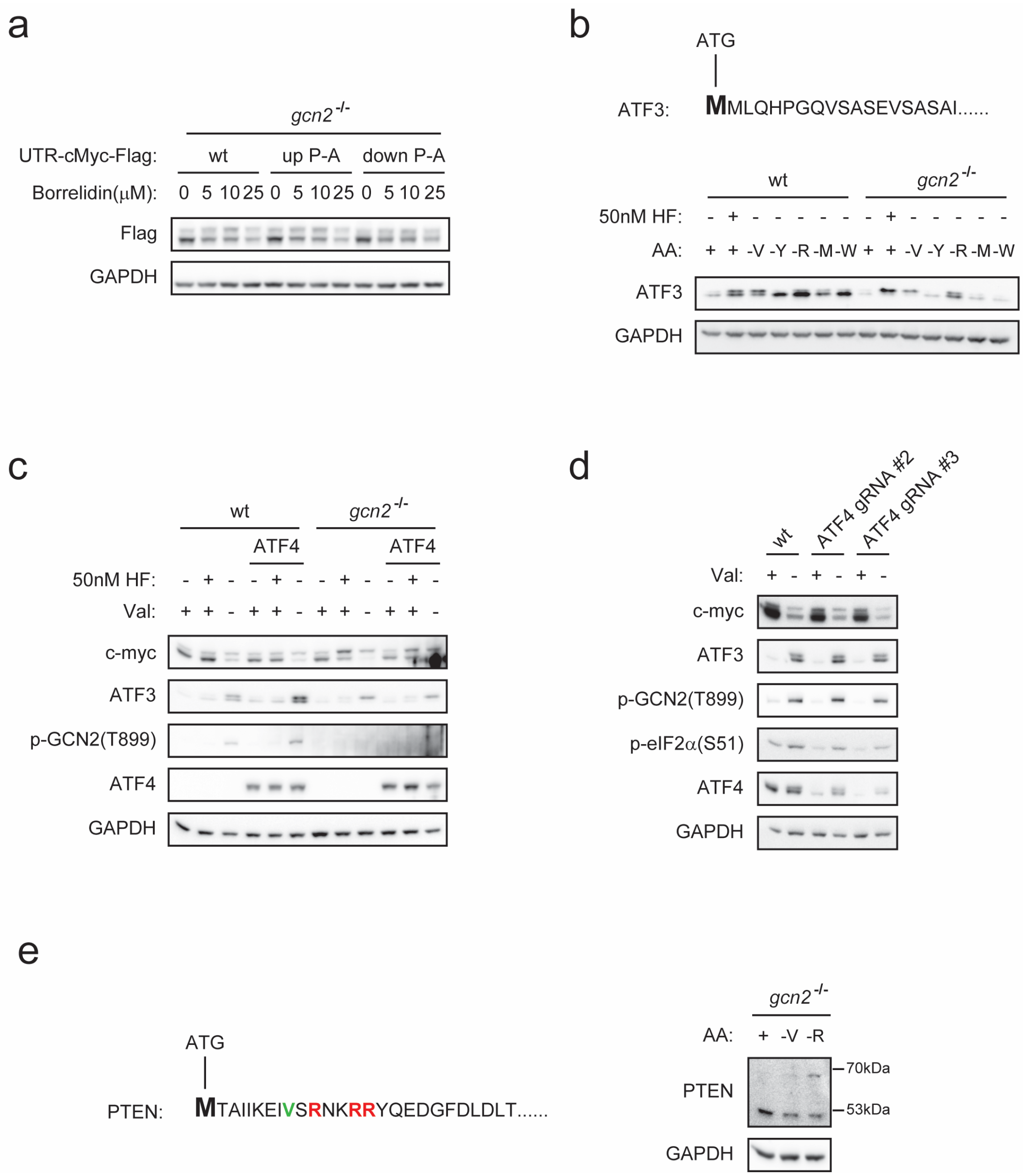


**Fig.S5**. **Effects of restriction of different amino acids on induction of distinct isoforms of c-myc, ATF-3 and PTEN.** a) HCT116(GCN2$^{-/-}$) cells were transfected with UTR-cMyc-Flag, UTR-cMyc$^{up\ P\text{-}A}$-Flag, UTR-cMyc$^{down\ P\text{-}A}$-Flag and treated with Borrelidin for 8 hours. b) HCT116$^{wt}$ or HCT116(GCN2$^{-/-}$) cells were treated with HF in standard DMEM, or incubated with DMEM medium without one of valine, tyrosine, arginine, methionine, or tryptophan for 8 hours. c) HCT116$^{wt}$ or HCT116(GCN2$^{-/-}$) cells were stably transformed with cDNA encoding human ATF4 and compared to untransformed controls with respect to c-myc-1 induction by HF or following incubation with DMEM without valine for 8 hours. d) HCT116$^{wt}$ or HCT116 cells in which ATF4 was inactivated by CRISPR (ATF4 gRNA#2,ATF4 gRNA#3) were incubated with DMEM without valine for 8 hours and analyzed for c-myc-1 induction. e) HCT116(GCN2$^{-/-}$) cells treated with HF in standard DMEM, or incubated with DMEM medium without one of valine or arginine for 8 hours, and analyzed by Western blot for PTEN isoforms.
